# Combinatorial effector targeting (COMET) for transcriptional modulation and locus-specific biochemistry

**DOI:** 10.1101/2024.10.28.620517

**Authors:** Caroline M. Wilson, Greg C. Pommier, Daniel D. Richman, Nicholas Sambold, Jeffrey A. Hussmann, Jonathan S. Weissman, Luke A. Gilbert

**Affiliations:** Tetrad Graduate Program, University of California, San Francisco, CA 94158, USA; Department of Urology, University of California, San Francisco, CA 94158, USA; Helen Diller Family Comprehensive Cancer Center, University of California, San Francisco, San Francisco, CA 94158, USA; Arc Institute, Palo Alto, CA 94304, USA; Johns Hopkins University School of Medicine, Baltimore, MD 21205, USA; Department of Computer Science, Stanford University, Stanford, CA 94305, USA; Prime Medicine, Cambridge, MA 02142, USA; Howard Hughes Medical Institute, Massachusetts Institute of Technology, Cambridge, MA 02139, USA; Whitehead Institute for Biomedical Research, Massachusetts Institute of Technology, Cambridge, MA 02142, USA; David H. Koch Institute for Integrative Cancer Research, Massachusetts Institute of Technology, Cambridge, MA 02139, USA; Department of Biology, Massachusetts Institute of Technology, Cambridge, MA 02139, USA

## Abstract

Understanding how human gene expression is coordinately regulated by functional units of proteins across the genome remains a major biological goal. Here, we present COMET, a high-throughput screening platform for <u>com</u>binatorial <u>e</u>ffector targeting for the identification of transcriptional modulators. We generate libraries of combinatorial dCas9-based fusion proteins, containing two to six effector domains, allowing us to systematically investigate more than 110,000 combinations of effector proteins at endogenous human loci for their influence on transcription. Importantly, we keep full proteins or domains intact, maintaining catalytic cores and surfaces for protein-protein interactions. We observe more than 5800 significant hits that modulate transcription, we demonstrate cell type specific transcriptional modulation, and we further investigate epistatic relationships between our effector combinations. We validate unexpected combinations as synergistic or buffering, emphasizing COMET as both a method for transcriptional effector discovery, and as a functional genomics tool for identifying novel domain interactions and directing locus-specific biochemistry.

## INTRODUCTION

Programmable DNA binding proteins have been widely leveraged to modulate gene expression and function across biology^1^. Fusion of diverse natural or synthetic domains to programmable DNA binding proteins alone or in combination has yielded artificial transcription factors, base editors, epigenome editors, modulators of 3D chromatin architecture, and much more^2–5^.

Synthetic CRISPR-based technologies enable robust, reversible, and multiplexable activation and inactivation of transcription when fused to epigenetic effector domains. Specifically, we and others have built catalytically inactive (dead) Cas9 (dCas9) fusion protein systems for transcriptional (CRISPRi and CRISPRa^6^) and epigenetic editing (CRISPRoff and CRISPRon^7^), enabling investigations of human gene function through genome-wide gain- and loss-of-functions perturbational experiments. In particular, CRISPRoff is a programmable epigenetic memory writer fusion protein, consisting of 4 domains: (1) a non-natural fusion of truncated domains from two DNA methyltransferase genes (DNMT3A-3L), (2) dCas9, and (3) a Krüppel-associated box (KRAB) domain^7^. CRISPRoff deposits both DNA methylation and repressive histone marks (H3K9me3) at the target genomic locus, and the combination of these activities results in synergistic and durable gene silencing that is not observed when each domain is fused to dCas9 individually. Given the complex interplay and cooperation between transcriptional modulators and epigenetic factors in human cells, CRISPRoff highlights how new technologies and biological properties can emerge by combining functional domains.

Beyond these individual rationally designed tools, much less emphasis has been placed on utilizing the CRISPR systems’ programmable DNA binding capabilities to scalably characterize the biochemical function of human proteins alone or in combinations when targeted to endogenous genes. Most existing approaches assay the effects of targeting individual proteins, protein domains or short sequences of proteins (∼80 amino acid tiles) to artificial fluorescent reporter genes^8^. Two recent efforts have evaluated the effects of pared tile combinations to artificial reporter genes and, separately, targeted individual protein tiles to endogenous and reporter genes to investigate context specificity^9,10^. Protein tiles although valuable for defining a minimal number of amino acids with some measure of biochemical effect are not physiologically relevant biophysical entities. To date, no approach has targeted combinations of proteins or full domains to endogenous genes to examine the combinatorial interactions between gene regulatory proteins that govern gene expression in natural systems at scale.

Here we present COMET, a platform for <u>com</u>binatorial <u>e</u>ffector targeting. We generate two libraries of combinatorial dCas9-based fusion proteins, containing one to three effector domains per terminus of dCas9, allowing us to use a pooled library functional genomics platform with a custom long-read sequencing readout to systematically investigate more than 110,000 combinations of effector proteins at endogenous loci for their influence on transcriptional activation or repression. Our fusion protein libraries incorporate protein domains from many areas of genome regulation, including writers and erasers of histone methyl and acetyl marks, chromatin remodelers, and DNA methyltransferases, in addition to positive control effector domains such as KRAB and VPR previously described in the literature^11–13^.

We observe more than 5800 significant hits (fusion proteins) that modulate transcription from our large-scale library screen in K562 cells and further implement our platform in induced pluripotent stem cells (iPSCs), suggesting that COMET can be implemented in alternative cell systems. Transcription factors, of which there are an estimated 1639 in the human proteome^14^, interact in an exquisitely complex fashion to coordinate gene regulatory networks. The ability of transcription factors to function independently or as concerted complexes dictates cellular processes, cell type specification, and ultimately the interpretation of DNA sequence^15,16^. Interestingly, in these experiments we identified far more unique transcriptional modulatory fusion protein hits than the total number of known transcription factors in the human proteome, underscoring the synergy in gene expression modulation that can arise in combinatorial space and drawing parallels to well-established principles of transcription factor cooperativity observed in vivo and in vitro^17,18^.

Further, we investigate epistatic protein relationships between our domain combinations that nominate unexpected biological protein relationships. We uncover novel synergistic domain combinations including a fusion protein containing 200 amino acids of methyl-CpG binding domain protein 2 (MBD2) and full-length chromodomain Y like (CDYL) protein that robustly represses transcription. Targeting MBD2 and CDYL together to a locus results in complete depletion of H3K27ac marks, relative to only partial loss of H3K27ac with either effector alone. Our domain epistasis analyses emphasize COMET as not only a platform for transcriptional effector discovery, but also as a functional genomics tool for identifying novel functional interactions between protein domains and for directing locus-specific biochemistry.

To make our COMET screening data on transcriptional modulation an accessible resource, we developed an interactive tool to visualize and explore our results (https://comet-ivp.gilbertlab.arcinstitute.org/). Overall, we propose COMET as a platform to directly probe functional effects of domain combinations on transcription at locus resolution.

## RESULTS

### A proof-of-concept combinatorial transcriptional effector library for modulation of endogenous human gene expression

We sought to design a functional genomics approach composed of a library of dCas9-guided transcriptional effector fusion proteins which can be used in a pooled screening assay to identify optimal domain combinations for transcriptional activation and repression at unperturbed endogenous human genomic loci. We designed two libraries that contain protein domains fused to both the N- and C-terminus of dCas9, rendering our synthetic fusion proteins RNA-guided (Figure 1A). Domains on either terminus of dCas9 are separated by linkers that have been engineered to reduce proteolysis and further, are long and flexible relative to the size of dCas9 (Figure 1B), such that we suspect our N- and C-terminal effectors to be able to interact in cis with each other or in trans with other human proteins. Elements in our libraries were selected to include protein domains (but excluding DNA binding domains) from known epigenetic modifier genes, as well as domains from previously characterized gene modulators^6,7,13,19^. Importantly, in our libraries we keep full length proteins or protein domains intact, allowing us to interrogate how entire protein domains or enzyme cores function in combination. As such, effectors in our libraries range from 138bp to 3870 bp, with a mean size of 1489 bp (median = 1305 bp), resulting in full length fusion proteins ranging from 4743-12198 bp (Figure 1C). Our small proof-of-method library consists of 37 protein domains fused on either terminus of dCas9, resulting in 1,369 dual terminus fusion proteins, hereby referred to as library 1 (L1) (Figure 1A and Supplementary Table 3). Elements in L1 include members of the Polycomb repressive complex 2 (PRC2), lysine methyltransferases and demethylases, deacetylases, and previously characterized gene modulators (VPR, DNMT3A, KRAB (CRISPRi), p300)^7,12,13,19^. Notably, a subset of these domains or full-length proteins are themselves composed of multiple domains, resulting in a total of 2-6 domains per dCas9 library element. For example, the VPR domain is composed of VP64, p65 and Rta activation domains.

**Figure 1.**
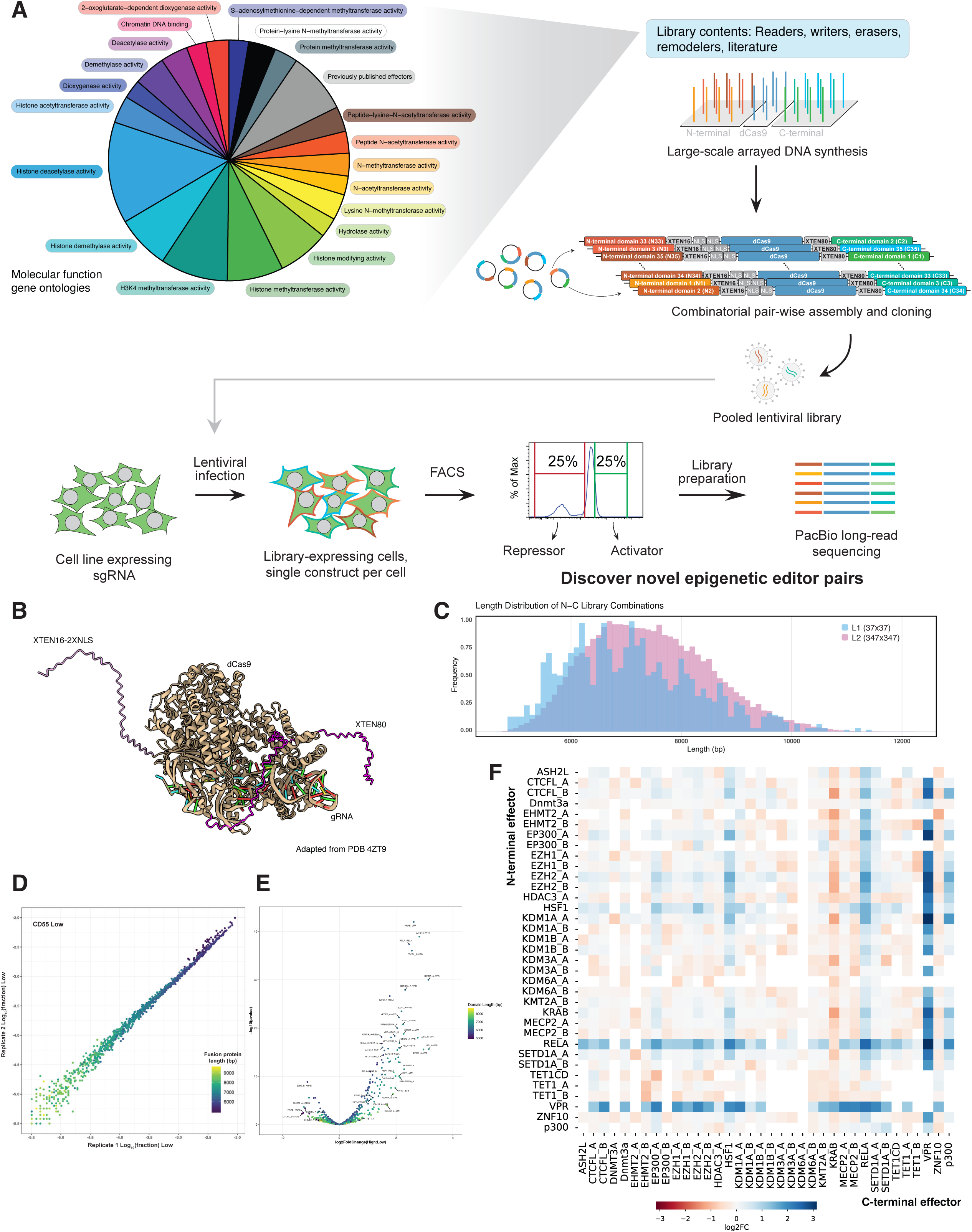
COMET generates combinatorial dCas9-based fusion proteins for endogenous gene targeting. (A) Pie chart showing library components as characterized by PANTHER GO-Slim gene ontology molecular signatures, showing categories containing ≥ 25 proteins, in addition to 81 effectors from the literature. Library elements were synthesized in an arrayed fashion, recoded for N- and C-termini of dCas9 and cloned in a pair-wise fashion. COMET screening schema showing pooled lentiviral library generation followed by infection into a gRNA-expressing cell line. Later in the screen, cells were harvested, stained for the target gene of interest, and binned by target gene expression for FACS. The top & bottom 25% of CD55 expressing cells were collected for library preparation and PacBio long read sequencing. (B) Structure of dCas9 in complex with a gRNA (PDB ID: 4ZT9^47^) with appended XTEN16-2XNLS linker (light purple) on the N-terminus of dCas9 and an XTEN80 linker (dark purple) on the C-terminus of dCas9, highlighting the long linker lengths with respect to dCas9’s 3D structure. (C) Histogram showing fusion protein lengths (bp) in the L1 (blue) and L2 (magenta) libraries. The length of dCas9 and the flanking linkers is 4464bp. (D) L1 COMET screen in K562 cells replicate correlation for the CD55 low condition showing the log_10_(fraction) of reads per library combination [Pearson correlation coefficient (R) CD55^low^ = 0.99 (p<2.2e-16)]. Points are colored by fusion protein length (bp). See also Supplementary Figure 1B. (E) DESeq of L1 screen data showing Log_2_Fold Change comparing the CD55^high^ to CD55^low^ conditions, with -log_10_(p-value) on the y-axis. Points are labeled with N- and C-domain names and colored by full length fusion protein length (bp). See COMET interactive volcano plots for further data visualization: https://comet-ivp.gilbertlab.arcinstitute.org/. (F) Heatmap of fusion protein combinations showing DESeq Log2FC values (activating in blue; repressive in red) with C-termini domains on the x-axis and N-termini domains on the y-axis.

We targeted *CD55* for transcriptional modulation using L1 in K562 cells, a myeloid cell model. CD55 encodes decay-accelerating factor, which plays an important role in inhibiting early complement activation^20^ and has been further implicated in numerous pathologies including malaria^21^, cancers^22^, and paroxysmal nocturnal hemoglobinuria^23^. Importantly, CD55 is expressed in K562 cells (Average log_2_TPM = 6.32)^24^ and previous work using CRISPR-based tools has demonstrated that we can both transcriptionally activate and repress *CD55* without impacting cell growth or survival^25^, allowing us to screen for both gene activation and repression phenotypes at a single endogenous locus.

We transduced L1 into K562 cells expressing a single gRNA targeting *CD55*. 8 days post-infection we stained for CD55 and sorted cells via FACS based on CD55 expression profiles (CD55-high and -low populations, along with an unsorted population). Full length effector constructs were amplified from genomic DNA and sequenced using PacBio long read sequencing (Supplementary Figure 1A). 92.8-93.4% of L1 was successfully represented in the screen (1270-1278 of 1369 possible combinations, replicate dependent) in the unsorted population of infected cells, with slight drop out of library elements that were not recoded for both termini prior to synthesis. We observed a high degree of correlation between replicates for each of the 3 conditions [Pearson correlation coefficient (R): CD55^high^ =0.99 (p<2.2e-16), CD55^low^ = 0.99 (p<2.2e-16), unsorted =0.98 (p<2.2e-16)] (Figure 1D and Supplementary Figure 1B-C), suggesting that our combinatorial transcriptional effector platform is robust and reproducible across replicates. To identify dCas9 fusion proteins that modulate CD55 transcription, we compared the CD55-high and -low samples to each other, generating enrichment scores for domains [Log_2_Fold Change (High:Low)].

In this initial proof-of-method L1 COMET screen, we analyzed our count data by testing for differential representation of each effector library element across CD55-expression conditions using DESeq2^26^, which nominated 187 enriched combinatorial effector hits out of 1007 total fusion proteins that were detected after count filtering (18.57%, p ≤ 0.05; Figure 1E-F). Significant hits (p ≤ 0.05) included both activators and repressors of *CD55*. We observed 146 significant activator combinations and 41 significant repressor combinations, emphasizing the strength of activation domains present in the proof-of-concept library. In a second analytic approach, we considered a combination enriched as an activator or repressor if it is represented greater than 2-fold in one population (high or low) relative to the other [abs(Log_2_Fold Change)>1], resulting in identification of both significant activators (n= 91) and repressors (n=8) in our L1 screen (Figure 1E). As further proof-of-method, known dCas9-guided transcriptional regulators including dCas9-KRAB, a robust transcriptional repressor used in genome-wide CRISPRi screens^6,27^, and transcriptional activator VPR^13^ (VP64-p65-Rta)-containing effectors were enriched in the CD55^low^ and CD55^high^ bins, respectively (Figure 1E), suggesting our platform can detect fusion proteins that can modulate CD55 expression in both directions. We observe unexpected repressive activities for specific domain combinations containing EHMT2_B and TET1 (both TET1_A and _B), where EHMT2 (also known as G9a) is a histone methyltransferase enzyme that writes mono- and dimethylated H3 Lys9 (H3K9me/me2)^28^, a hallmark of heterochromatin, and TET1 is an DNA demethylase that oxidizes 5-methylcytosine resulting in gene activation^29^ (Figure 1F). Notably, we observe that VPR, a tripartite transcriptional activator used for gene activation functional genomics applications, is enriched in the CD55^high^ bin even when fused to known transcriptional repressors like KRAB or MeCP2^30^, suggesting that our screening platform can be used to uncover novel domain epistasis phenomena wherein the activity of one domain supersedes the activity of a second domain (Figure 1F).

### Large-scale combinatorial transcriptional effector discovery enables robust endogenous human gene modulation

To build on our COMET platform for combinatorial effector discovery, we generated a second larger-scale library (L2) consisting of 347 effector domains fused in all pair-wise combinations to the N or C-terminus of dCas9 (Figure 1A & Supplementary Table 3). 266 of these domains were isolated from 194 proteins nominated by the dbEM, a database of epigenetic modifiers^31^. 81 previously published effectors were included as positive controls (Supplementary Table 1). The small-scale library elements are encompassed in the large-scale library as an internal control. Of the 347 domains in the library, 110,554 combinations of a possible 120,409 combinations were synthesized and cloned successfully after domain synthesis dropout (Figure 1C).

We performed a large-scale effector discovery screen in K562 cells in which L2 proteins were recruited to the target locus, *CD55*, via a single gRNA (as described in the L1 screen, Figure 1A). Given our use of PacBio long-read sequencing, our sequencing depth was limited for two replicates of both high- and low CD55-expressing cells, such that 72,547-77,472 of 110,554 domains were detected (65.6-70.1%, replicate and condition specific) prior to count threshold filtering. After filtering for fusion proteins with 10 counts per replicate and condition, counts were normalized and Log_2_Fold Change (High:Low) was calculated (see methods). Despite low library sequencing coverage, our results from the L2 screen were highly correlated across replicates for both CD55^high^ and CD55^low^ conditions [Pearson correlation coefficient (R): CD55^high^ =0.89, p-value < 2.2e-16; CD55^low^ = 0.92, p-value < 2.2e-16] (Figure 2A and Supplementary Figure 2A). Consistent with observations using L1, we observed enrichment of positive control domains including KRAB and VPR in CD55^low^ and CD55^high^ bins, respectively (Figure 2B). We observed 5805 significant hits (p-value <= 0.05); 2337 (40.3%) increased CD55 expression more than 2-fold, and 2074 (34.7%) repressed CD55 expression more than 2-fold (Figure 2B-C and Supplementary Figure 2B). Of the 5805 significant combinations nominated in our L2 COMET screen, 98.9% (5744) are fusion proteins that have effector lengths (N + C) greater than 480bp–the equivalent of two 80 amino acid tiles fused to dCas9, underscoring full length domain combinations as a rich resource for transcriptional perturbations (Figure 2D).

**Figure 2.**
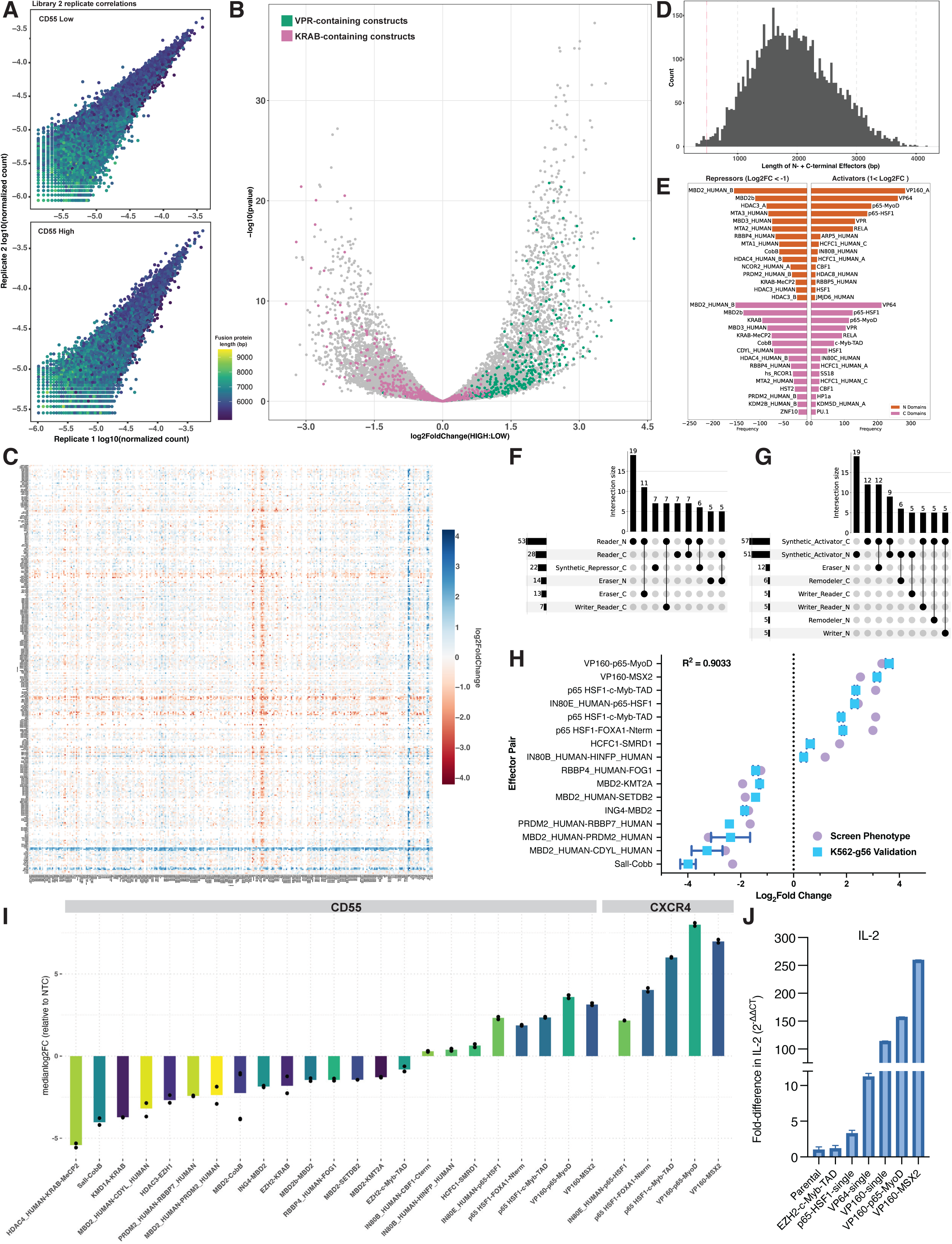
Large-scale combinatorial transcriptional effector discovery enables robust gene modulation. (A) L2 COMET screening platform replicates are highly correlated. Plots show CD55^low^ (top) and CD55^high^ (bottom) replicate correlations using normalized counts (fraction) for domain combinations. Pearson correlation coefficient (R): CD55^high^ =0.89, p-value < 2.2e-16; CD55^low^ = 0.92, p-value < 2.2e-16. Points are colored by fusion protein length (bp). See also Supplementary Figure 2A. (B) DESeq for L2 screen in K562 cells showing Log_2_FC (High:Low CD55 expression bins) on the x-axis and -log_10_(p-value) on the y-axis, with 5805 significant fusion protein combinations (p-value ≤ 0.05). See COMET interactive volcano plots at https://comet-ivp.gilbertlab.arcinstitute.org/. (C) Heatmap of Log_2_FC (High:Low CD55 expression bins) for individual N-C effector combinations. N domains are on the y axis and C domains are on the x axis, with positive log2FC in blue (activators) and negative log2FC values in red (repressors). See also Supplementary Figure 2B for enlarged format. (D) Histogram of 5085 significant L2 screen effector lengths (N-terminal + C-terminal, excluding constant dCas9 sequence of 4464bp), showing 5744 domain pairs longer than 480 bp total (pink dashed line). (E) Top 15 most frequently observed significant (p-value < 0.05) N or C domains for repressors (left, log2FC < -1) and activators (right, log2FC > 1) from the L2 K562 screen at CD55, showing enriched N effectors in orange and C effectors in pink. Frequency is shown on the x-axis. See also Supplementary Figure 3A. (F&G) Upset plots categorizing N and C domain function for the 100 strongest (F) repressors (log2FC < -1) and (G) activators (log2FC > 1) from the L2 K562 screen with an intersection size ≥5. See also Supplementary Figure 3B-C. (H) Screen phenotype (log2FC) values from DESeq for individual effector combinations (purple circle) compared to arrayed validation (blue square, 2 replicates) gene expression fold change values as measured by flow cytometry relative to a dCas9 protein with inactive N- and C-terminal domains (pCW31) on day 3 post-transduction. Screen and arrayed validation log2FC values are highly correlated [Pearson R squared: 0.9034, p-value (two-tailed) <0.001]. (I) Arrayed validation of COMET nominated constructs in dual-gRNA expressing K562 cells targeting the *CD55* locus or *CXCR4* locus as measured by antibody staining and flow cytometry 3 days post-transduction. Bar plot values represent the median log2Fold Change of target gene expression (CD55 or CXCR4) per effector combination normalized to the median target gene expression in cells transduced with negative control dCas9 protein (pCW31) for two replicates per transduction. Cells are gated on stable gRNA expression (BFPpos), followed by mNeonGreen fluorescence as a proxy for transduction efficiency/protein expression. (J) Fold change expression of IL-2 as measured by qPCR in K562 cells expressing a dual-gRNA targeting *IL-2* and transduced with activator constructs. IL2 expression is normalized to guide-expressing parental K562 cells and GAPDH is the endogenous control. Mean CT values (run in triplicate) and standard deviations are used in the ΔΔCT calculations.

Of the significant repressor combinations nominated in the L2 screen (log2FC < -1 & p-value < 0.05), the most frequently represented individual effector domains included domains from the methyl-CpG-binding domain (MBD) protein family including MBD2, MBD2b, MBD3, and MeCP2, as well as nuclear coreceptor co-repressor 2 (NCOR2) and some of its histone deacetylase interaction partners HDAC3 and HDAC4 (Figure 2E & Supplementary Figure 3A). The presence of HDACs as strong repressors within our screen emphasizes the potential use case of chromatin context-specific editors, such that markers of active gene expression at defined loci including H3K27ac could be targeted for removal and further repressed by a second transcriptional repressor. Unexpectedly, we observe chromodomain Y-like protein (CDYL) as a frequent C-terminal repressor (Figure 2E). CDYL is a protein known to bind repressive chromatin marks including H3K9me2/3 and H3K27me3^32,33^, that is relatively understudied and has not previously been fused to dCas9 for direct transcriptional perturbations. On the activation side (1 < log2FC & p-value < 0.05), we most frequently observe strong acidic transactivation domains and synthetic activators including VP64, VPR, p65-HSF1 (Figure 2E & Supplementary Figure 3A). Unexpectedly, we also observe effectors that are not commonly considered for dCas9-based activation approaches including viral accessory protein host cell factor C1 (HCFC1) and INO80 family subunits (Figure 2E).

We also investigated what types of domains most frequently functioned together as repressors or activators. We categorized our library elements by defining whether proteins contained domains that function as chromatin remodelers, or readers, writers, or erasers of DNA methylation, histone methylation, or histone acetylation, using preexisting classifications^34^ and InterPro annotations^35^. For example, of the 100 strongest repressors we observed, 53 N-terminal domains were from proteins with known reader function (Figure 2F and Supplementary Figure 3B), where reader domains are thought to engage with an epigenetic mark (or many) and modulate transcription, chromatin organization, and/or DNA repair. Further, 11 of the 100 strongest repressors included an N-terminal reader and C-terminal eraser, highlighting the potential for synergy between the deposition of a repressive mark and the erasing of an activating transcriptional mark (Figure 2F and Supplementary Figure 3B). Of our strongest activators (top 100), synthetic activators were present on either the N or C terminus of the fusion protein in 57 & 51 combinations, respectively (Figure 2G and Supplementary Figure 3C). Notably, with the inclusion of additional domains in L2 relative to L1, we see an increase in the range of tunable CD55 expression, where log_2_FC values ranged from -1.3 to 3.14 using L1, compared to -3.44 to 4.2 in the L2 K562 screen (Figure 1E & 2B), emphasizing our system as a platform for epigenetic editor discovery and tunable locus-specific gene expression depending on the domain pair combination. Our COMET screening data is interactively explorable at https://comet-ivp.gilbertlab.arcinstitute.org/.

To validate our L1 and L2 screening results and to begin to explore the combinatorial biology that emerges from these experiments, we compared screen phenotypes (Log_2_FC enrichment scores) to measurements of gene activation or repression in a validation context, using the same single gRNA targeting *CD55* as that of the screens. In an arrayed format, we targeted *CD55* with 16 different effectors, along with an inactive dCas9 control. Our results show strong correlation of effector activity across both screen and arrayed experiments (R^2^ = 0.9033) suggesting that our domain screening platform produces reproducible enrichment scores that translate to functional gene modulation effects (Figure 2H).

Confident in our platform for nominating novel activator and repressor combinations, we cloned several additional hit fusion proteins for arrayed validation and evaluated their functionality at *CD55*, *CXCR4*, and *IL-2*, to demonstrate hit proteins are active at more than one endogenous human gene. We delivered effector constructs of interest via lentivirus to K562 cells stably expressing a dual gRNA construct targeting the indicated endogenous gene (Supplementary Table 2) and then measured target gene expression levels between 3 to 9 days post-transduction via antibody staining and flow cytometry for surface proteins. At *CD55*, we observed robust repression or activation on day 3 post-transduction relative to a scrambled negative control protein (Figure 2I). CD55 targeting by COMET effectors resulted in a >40-fold decrease in expression (HDAC4-KRAB-MeCP2) to a >11-fold increase in CD55 expression (VP160-p65-MyoD), suggesting our library can identify effectors in a large dynamic range of gene modulatory activity. To evaluate the generalizability of our platform in contexts in which the target gene is off or not appreciably expressed, we targeted *CXCR4* and *IL-2*. We observe robust activation of CXCR4 measured by flow cytometry ranging from >4.6 fold increase in gene expression (IN80E-p65-HSF1) to more than 240-fold increase (VP160-p65-MyoD) relative to a negative control (scrambled) dCas9 (Figure 2I, right). Further, we observe *IL-2* activation in K562 cells as measured by qPCR, with the strongest activation from effectors nominated by our domain screening platforms compared to known activators including p65-HSF1 and VP64/160, suggesting that our platform can identify highly effective fusion protein combinations that can activate genes from an off-state (Figure 2J). Further, our effectors remain active at later timepoints (day 6 post-transduction), suggesting that they are neither silenced nor toxic to K562s despite constitutive expression (Supplementary Figure 2C). Importantly, COMET effectors have a wide dynamic range of gene expression modulation, enabling systematic investigations of how tunable and targeted gene dosage influences cellular function and genetic interactions.

### Combinatorial effector discovery in iPSCs

It is widely known that the efficacy of CRISPR-based effectors can vary in different cell types, due to effector toxicity, protein dosage limitations, and variations in transgene silencing mechanisms^36^. As such, we tested the efficacy of COMET effectors in KOLF2.1J iPSCs. We performed a domain library screen as described above in iPSCs at *CD55* using L1 and the same guide as that of the K562 screens, in order to better understand cell-type specific effector combinations. iPSC screen replicates were highly correlated [Pearson correlation coefficient (R): CD55^high^ =0.99 (p<2.2e-16), CD55^low^ = 0.99 (p<2.2e-16)] (Figure 3A). We observe 42 significant hits from our stem cell screen, 30 of which were unique to the iPSC screen and not significant in our L1 screen in K562 cells (Figure 3B). Further, we observe similar dynamic range of transcriptional modulation as that of L1 in K562 cells, suggesting that our platform can be readily translated to alternative cell types.

**Figure 3.**
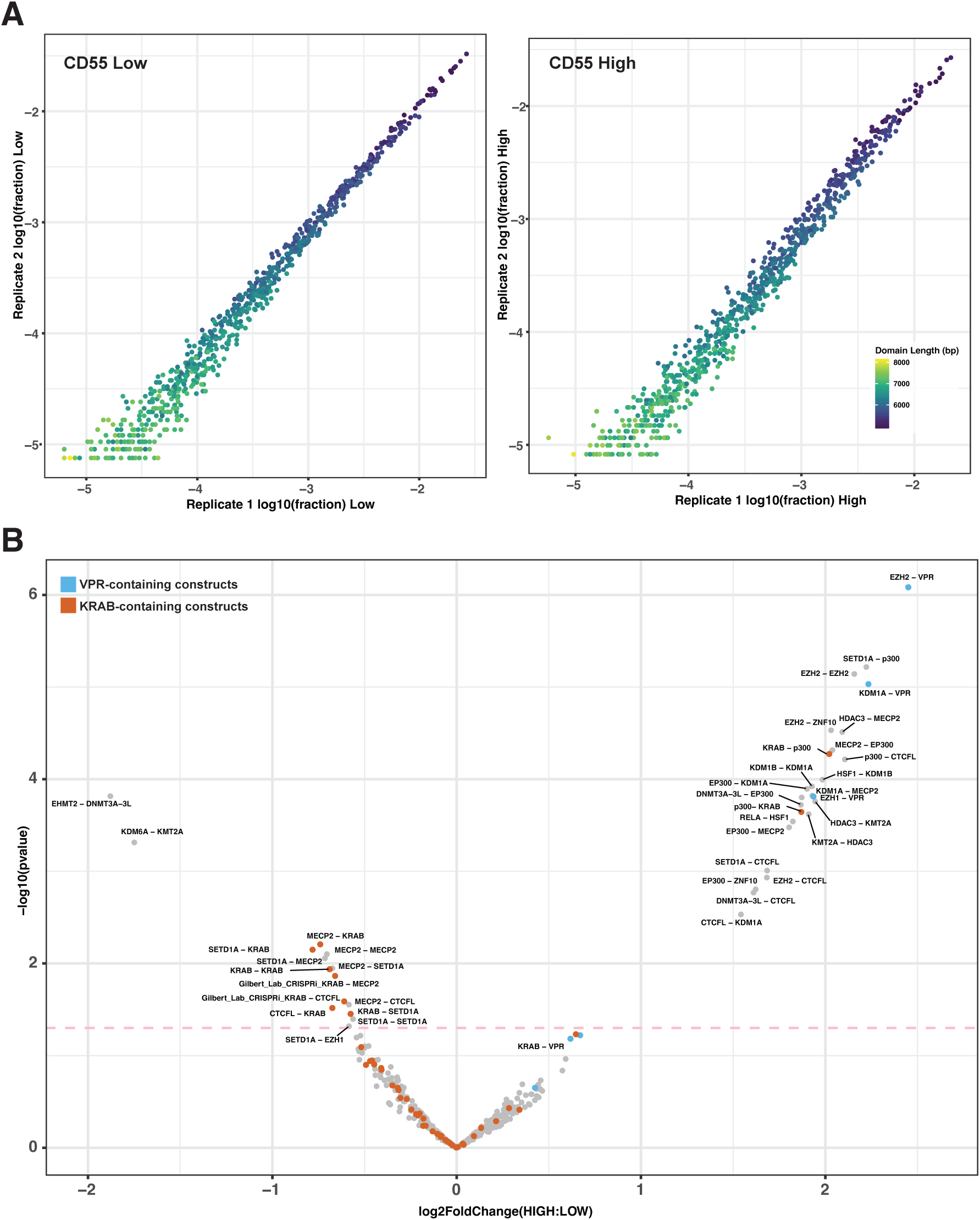
Combinatorial effector discovery in iPSCs. (A) Replicate correlations for COMET L1 screen in KOLF2.1J iPSCs targeting the *CD55* locus, showing log_10_(fraction) of reads per effector combination for CD55^low^ and CD55^high^. Pearson correlation coefficient (R): CD55^low^ = 0.99 (p<2.2e-16), CD55^high^ =0.99 (p<2.2e-16). (B) DESeq for L1 screen in iPSCs showing Log_2_FC (High:Low CD55 expressing bins) on the x-axis and -log_10_(p-value) on the y-axis, with 41 significant fusion protein combinations (p-value ≤ 0.05). A count threshold of 5 was used for all conditions. See COMET interactive volcano plots at https://comet-ivp.gilbertlab.arcinstitute.org/.

### Combinatorial effector screening enables discovery of domain epistasis

Motivated by the synergy of repression observed between effector domains in our previous work developing CRISPRoff^7^, we sought to use COMET to identify unexpected effector combinations; wherein the N-C fusion protein phenotype deviates from the additive model derived from measuring the phenotypes of individual N or C terminus effectors alone (Figure 4A). Our library does not contain single effectors fused to dCas9, so we estimated single domain phenotypes by fitting a least squares regression for each library element. For each domain combination in our library, we calculated least-squares expected combination phenotypes based on the null assumption that the domains do not phenotypically interact. We compared this expected value to the observed combination enrichment scores from the L2 screen per replicate and utilized this model to identify unexpected domain-domain relationships, in which specific combinations led to phenotypes that were greater than or less than the expected phenotype (Figure 4B-C and Supplementary Figure 4A). We term this difference between the expected and the observed phenotype the “residual” value (Figure 4B-C and Supplementary Figure 4B). We observed, as expected, that most domain combinations (55.75%) have a score that is close to the expectation of no interaction between domains, in which the observed activity is within +/-0.3 of the predicted activity. Uncovering strong unexpected functional relationships between domains, however, is rare; only 3.52% of effector combinations in our epistasis analysis have residuals >1 or <= -1 (Figure 4C). Interestingly, the observation that most domains do not interact and that strong interactions are rare mirrors findings from studies on gene-gene interactions^37^. These results also demonstrate the COMET platform can scalably nominate protein domain combinations that act epistatically to modulate a transcriptional phenotype in living human cells.

**Figure 4.**
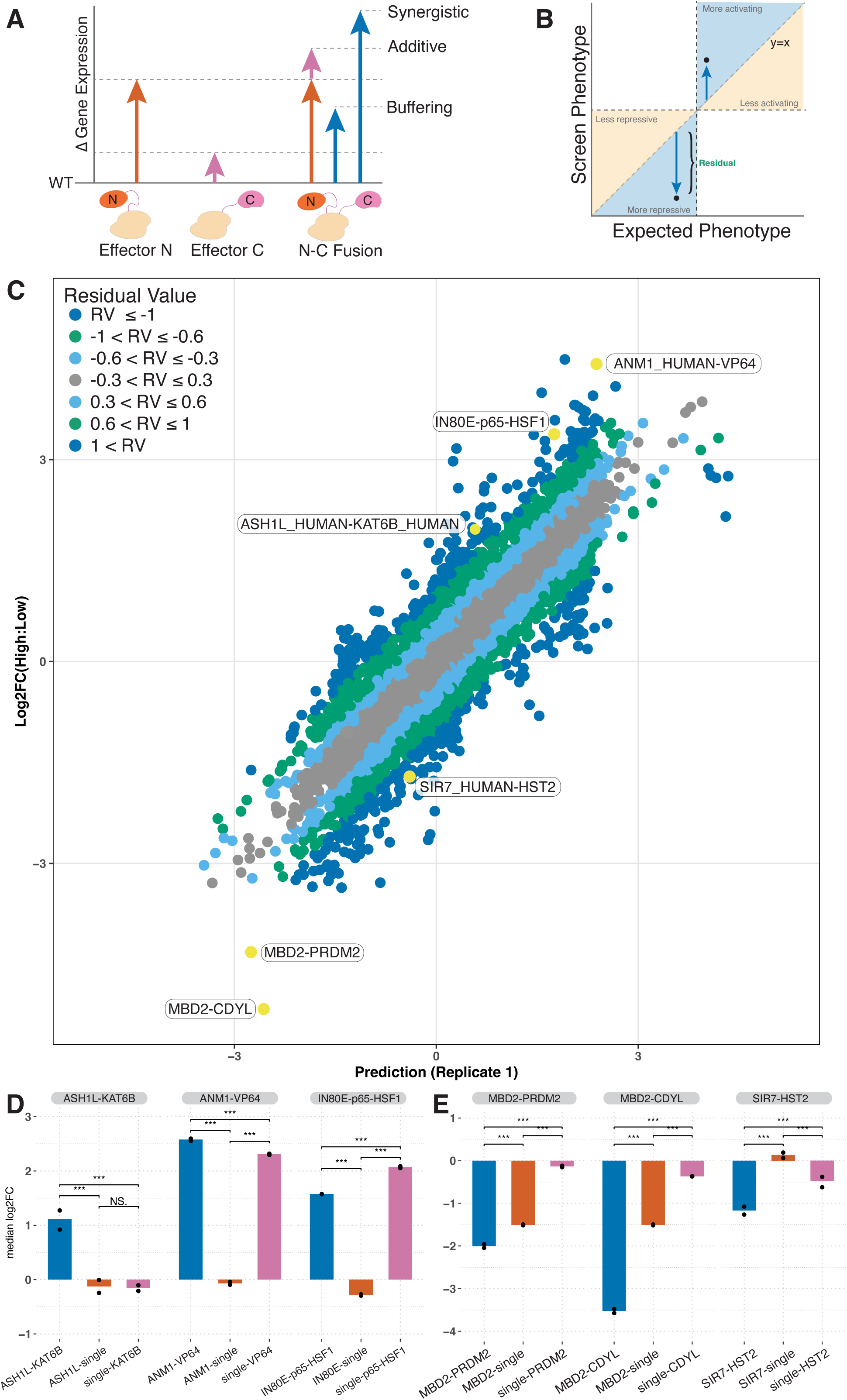
COMET screening enables discovery of domain epistasis. (A) Schematic of changes in gene expression depending on N-, C-, or dual fusion activity, wherein activity greater than the combination of individual activities is synergistic. N-C fusions with activity less than the activity of an individual effector is buffering or antagonistic. (B) Schematic of residual analysis comparing the expected phenotype to the observed screen phenotype, where the residual is the difference between our observed and expected phenotypes. (C) Scatter plot showing epistasis modeling predictions for effector combination activity on the x-axis for Replicate 1 and DESeq2 Log2FC values for effector combinations from the COMET L2 screen in K562 cells on the y-axis. Colored points represent zones of residual values, i.e. the distance between the expected vs observed gene modulation activity at the *CD55* locus. 29.23% of combinations have residuals between 0.3 and 0.6 or -0.3 and -0.6 (light blue); 11.50% of combinations have residuals between 0.6 and 1 or -0.6 and -1 (green); and 3.52% of combinations have residuals >1 or <= -1 (dark blue). 55.75% of combinations have minimal residuals (less than 0.3) (gray). Combinations nominated with residuals >1 (dark blue zone) were subsequently cloned as double and single-dCas9 effectors to evaluate synergy. (D & E) Arrayed validation of nominated effectors by flow cytometry, with median log2FC (normalized to an inactive dCas9 fusion protein) on the y-axis. Dual and single terminus fusion proteins were cloned and transduced into single CD55 gRNA-expressing K562 cells. (D) ASH1L_HUMAN (N172)-KAT6B_HUMAN (C106); ANM1-VP64; IN80E-p65-HSF1. (E) MBD2-PRDM2 (PRDM2_HUMAN_B, C131); MBD2-CDYL; and SIR7-HST2. Wilcoxon t-tests per replicate per comparison were performed in R (*** denotes p-value <2e-16).

To validate these findings, we cloned and tested residual-nominated constructs in an arrayed fashion for functional domain epistasis analysis. In these experiments, we compared the residual-hit protein to fusion proteins where the N- or C-terminus domains were replaced with an inactive scrambled peptide to generate effectors with a single active domain fused to dCas9. The activity of single domains were then compared to their respective double domain activity using the same single guide targeting *CD55* that was used in the screen. In our validation experiments, we tested epistatic domain interactions nominated as synergistic and antagonistic. We observe gene activation synergy with the C-terminal portion of ASH1L (also known as KMT2H) and the first 213 aa of KAT6B, a histone acetyltransferase, where each effector individually shows minimal changes in CD55 expression (Figure 4D). We also observe gene activation synergy with the effector combination of full length ANM1 (also known as protein arginine methyltransferase 1, PRMT1) and VP64, wherein ANM1 alone showed no modulation of CD55 expression and VP64 robustly activated CD55, but together, we observe even greater activation of CD55 (Figure 4D). We observe antagonistic effects within INO80E-p65-HSF1, the combination of a full length INO80 family chromatin remodeler and bipartite transcriptional activator^38^, in which the individual domain effect of INO80E negatively affects the activation observed with p65-HSF1 alone (Figure 4D). We observe modest synergy in CD55 repression by MBD2-PRDM2, which consists of the C-terminal portion of MBD2, a methyl-CpG-binding domain protein, and 627 amino acids of PRDM2, a histone/protein methyltransferase (Figure 4E). We observe synergy for SIR7-HST2, a combination of full-length human and *Saccharomyces cerevisiae* deacetylases, respectively, in which SIR7 alone is inactive at CD55, but in combination with HST2 results in a reduction in CD55 expression by more than 2 fold (Figure 4E). In the case of MBD2-CDYL, we observe repression of CD55 with MDB2 alone, and minor repression with CDYL alone, where CDYL is chromodomain Y-like protein (CDYL). However, when in combination, we observe greater than 11-fold knockdown of CD55 (Figure 4E), suggesting a synergistic mechanism of repression at the target locus. These experiments validate that COMET data can nominate unexpected epistatic relationships between protein domains that modulate transcriptional phenotypes.

We were intrigued by the strong repressive synergy between MBD2 and CDYL, as CDYL is poorly characterized and this combination of proteins has not been studied. As noted above, CDYL was identified as a common C-terminal repressor in our screen (Figure 2E), however most repressive combinations of domains are not measured to be synergistic, suggesting our COMET residual analysis enabled discovery of an interesting biological interaction (Figure 4C). MBD2 is known to bind methylated DNA; however, in our COMET fusion protein library we excluded DNA binding domains naturally present in human proteins. As such this MDB2 domain (N105 in our library) excludes the N-terminal MBD domain and includes the p55-binding region and the C-terminal domain (Figure 5A). CDYL has been implicated in interactions with G9a, a writer of H3K9me1/2, and in enhancing deposition of H3K27me3 by PRC2^32,33,39^. We sought to understand the mechanism of MBD2-CDYL repression. We hypothesized that MBD2-CDYL could be recruiting histone deacetylases to *CD55*, through interactions between MBD2 and the NuRD complex, or in CDYL’s interactions with HDAC1/2^40^. To test this, we performed CUT&Tag to probe histone 3 lysine 27 acetyl (H3K27ac) marks at the target locus and observed a near complete loss of H3K27ac at the *CD55* locus in cells targeted with the MBD2-CDYL effector relative to parental K562s. Interestingly, targeting by single effector domains (MBD2 and CDYL each fused to dCas9 alone) each individually resulted in a partial decrease in H3K27ac relative to parental K562s, but to a lesser extent than targeting with the combined MBD2-CDYL effector, suggesting that strong deacetylation is a product of MBD2-CDYL when combined at the target locus (Figure 5B & C). In independent repeat experiments assaying H3K27ac marks at the target locus by CUT&Tag, we observed a consistent decrease in H3K27ac when targeted by MBD2-CDYL relative to parental controls (Supplementary Figure 5A). By contrast, we do not observe noticeable deposition of H3K9me1 marks at the *CD55* locus as measured by CUT&Tag when targeted by MBD2-dCas9-CDYL, MBD2-dCas9 or dCas9-CDYL (Supplementary Figure 5B), suggesting that transcriptional repression induced by this protein combination in this context is not dependent on G9a or G9a-like protein (GLP) deposition of H3K9me1 repressive marks. To rule out alternative mechanisms of silencing via potential interactions between CDYL and members of PRC2, we also performed CUT&Tag for H3K27me3 and observe no deposition of H3K27me3 at the *CD55* locus when targeted by MBD2-dCas9-CDYL, MBD2-dCas9 or dCas9-CDYL relative to control cells (Supplementary Figure 5C). It is possible that in this context MBD2-dCas9-CDYL acts to repress transcription mainly by recruiting histone deacetylase complexes or it is possible additional repressive mechanisms are recruited by MBD2-CDYL.

**Figure 5.**
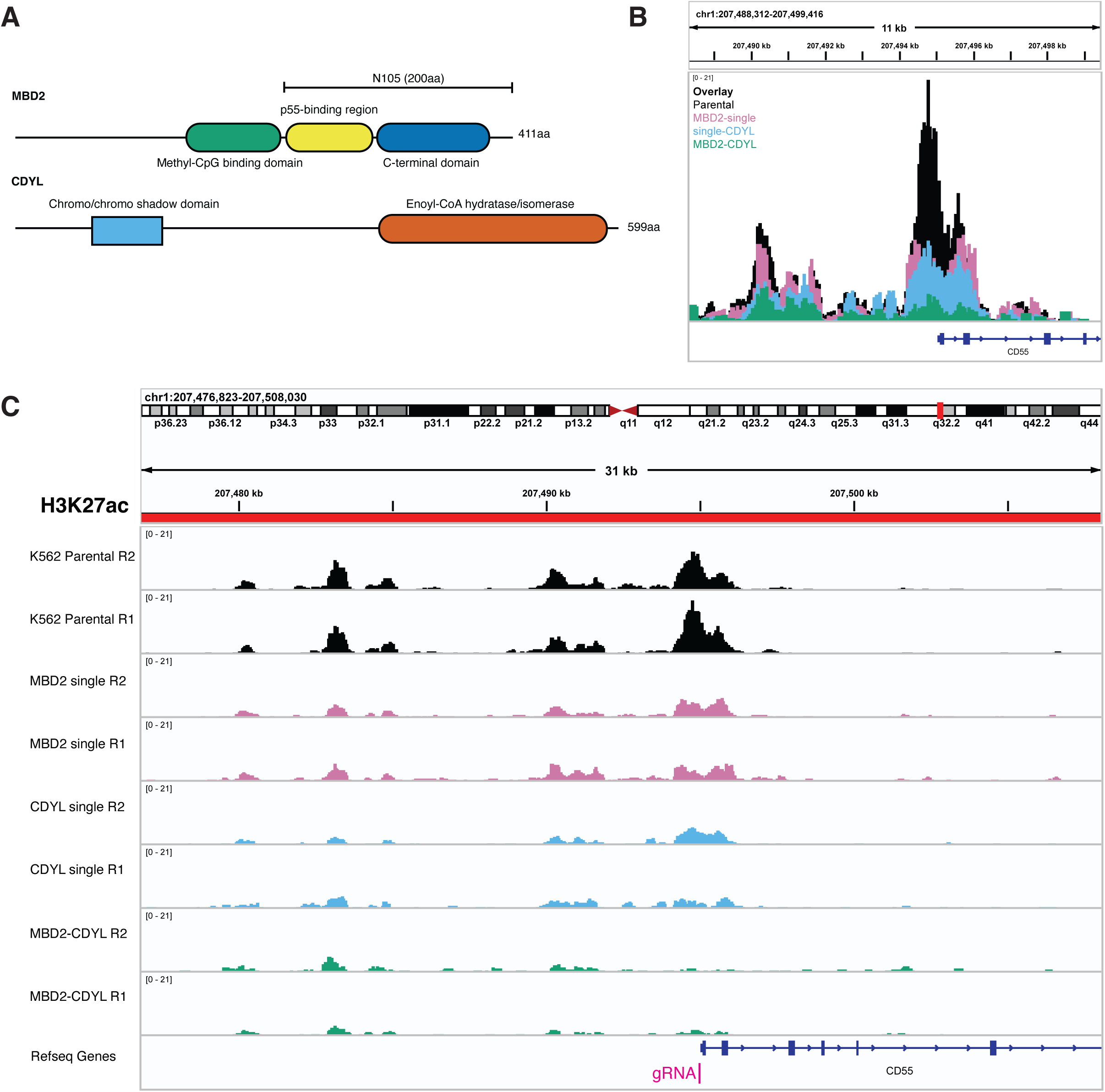
MBD2-CDYL exhibit repressive synergy and ablation of H3K27ac at *CD55* locus. (A) Schematic of MBD2 and CDYL domain architecture. The C-terminal region of MBD2 (200aa, shaded yellow box) is included in the specific effector combination we evaluated with full length CDYL (598aa). (B) Overlay of H3K27ac peaks in IGV for all 8 samples (parental K562s, MBD2-single, single-CDYL, and MBD2-CDYL, with 2 replicates) at an 11kb region around the transcription start site (TSS) of *CD55*. (C) H3K27ac CUT&Tag data for parental K562 cells (black), or cells transduced with MBD2-single (pink), single-CDYL (light blue), and MBD2-CDYL (green) effectors across two technical replicates as visualized in IGV across a 31kb region surrounding the TSS. Position of the guide RNA is denoted in pink.

Overall, our platform nominates novel combinatorial fusion proteins that exhibit a wide dynamic range of transcriptional activation and repression at endogenous loci. Through the use of full-length protein domains and flexible linkers, our fusion proteins allow for direct catalytic mechanisms of gene regulation and maintain protein surfaces for at-locus protein-protein interactions. We demonstrate that COMET can be implemented to nominate synergistic domain combinations, whose mechanisms suggest targeted writing or removal of biochemical marks in a locus-specific manner.

## DISCUSSION

Here, we present COMET, a combinatorial effector targeting platform for the discovery of novel transcriptional modulators. Through the use of full-length effector domains combinatorially fused to dCas9, we evaluate the activity of more than 110,000 domain combinations on transcription at endogenous human loci. We identify thousands of novel effector combinations for locus-specific gene activation or repression, and further, work to dissect unexpected combinations in our library.

It has been challenging to understand how the proteomic landscape and chromatin environment directly influence transcription at individual loci. This can be done in vitro to understand how enzymes and remodelers influence chromatin accessibility or epigenetic marks, but such approaches do not account for the dynamic chromatin landscape or how such effectors directly influence transcription in living human cells. Alternative approaches to understanding locus-specific environments utilize mass spectrometry (MS) following target amplification, dCas9-based enrichment strategies, or proximity labeling based approaches^41–44^. These enrichment strategies, however, have high background signal and more importantly are correlative by the nature of the assays. As such, we need better approaches for functionally defining how proteins, protein domains, or enzyme activities when localized to an individual endogenous human genomic loci modulate transcription. We propose COMET as a solution for this very purpose, as we can characterize the effects of thousands of effectors in a locus-specific fashion and evaluate their effects on transcription or potentially other nuclear phenotypes. Importantly, our results show we can write and erase biochemical marks at individual locations in the genome.

By including long flexible linkers on either terminus of dCas9, we suspect that effectors in our library can interact in cis or in trans to activate or repress loci. This approach enables discoveries of synergistic effector combinations and investigations of domain epistasis in targeted genomic contexts. Additionally, by keeping full-length domains and enzyme cores intact, we preserve catalytic sites and surfaces/scaffolds for protein-protein interactions. Of the >5800 significant hits in our large-scale library, the median length of both effectors (excluding dCas9 and linkers) was 632 amino acids, highlighting the utility of full-length effectors in producing robust activation and repression (Figure 2D). For example, in the case of our MBD2-CDYL synergistic effector, the MBD2 effector contains two annotated domains, both of which may play a role in the repression we observe. Specifically, our effector contains both the p55-binding domain and the full C-terminal domain, which is known to contain a coil involved in coiled-coil interactions between MBD2 and transcriptional repressor p66-alpha (GATAD2A), a histone deacetylase and component of the NuRD complex^45,46^. The inclusion of both the transcriptional repressive domain and the C-terminal domain of MBD2, which are 198 amino acids in length, would not have been possible with current 80 amino acid tiling approaches, and it is unlikely that we would have observed the same level of synergy with CDYL had we not used the full-length protein (599 amino acids). In conclusion, COMET enables scalable characterization of human proteins and protein domains that act alone or in combination to modulate transcription and these results enable us to quantitatively measure epistatic interactions between protein domains that lead to biochemical hypotheses on the nature of transcriptional control of human gene expression at specific genomic loci.

## ACKNOWLEDGEMENTS

The authors thank Aidan Winters, Sarah Hsu, Garret Wong, Chris Hsiung, James Nuñez, Fyodor Urnov, April Pawluk, Brian Plosky and members of the Gilbert lab for their thoughtful discussions and feedback. The authors thank Twist Bioscience and the Innovation Lab for their work generating the library elements. The authors thank Scott Nanda, Coco Wu, Vijay Ramani, and members of the Ramani lab for their work running the Sequel IIe and coordinating data transfers. The authors thank Nick Youngblut for his contributions to the COMET interactive volcano plot Shiny app. The authors thank Oanh Nguyen and the DNA Technologies and Expression Analysis Core at the UC Davis Genome Center, supported by NIH Shared Instrumentation Grant 1S10OD010786-01. The authors thank Sarah Elmes and members of the UCSF Helen Diller Family Comprehensive Cancer Center Laboratory for Cell Analysis (LCA), supported by NIH NCI Award Number P30CA082103.

This research was funded by the DARPA PREPARE program (HR00111920007 to L.A.G. and J.S.W.) and by the Arc Institute. L.A.G. is funded by the Arc Institute, NIH (DP2CA239597, UM1HG012660), CRUK/NIH (OT2CA278665 and CGCATF-2021/100006), and a Pew-Stewart Scholars for Cancer Research award.

## AUTHOR CONTRIBUTIONS

L.A.G., J.S.W., G.C.P., and C.M.W. conceived of the study and also designed and interpreted experiments. G.C.P. and L.A.G. designed the libraries of combinatorial fusion proteins. G.C.P. performed proof-of-concept implementation of the COMET platform. G.C.P. and C.M.W. performed library screens. J.H. wrote the long-read sequencing alignment pipeline. C.M.W. performed screen data analyses. C.M.W. cloned arrayed validation constructs with help from N.S. C.M.W. performed validation experiments. D.D.R. performed epistasis modeling. C.M.W. performed epistasis validation and CUT&Tag experiments. C.M.W. and L.A.G. wrote the manuscript with input from all authors.

## DECLARATION OF INTERESTS

C.M.W., G.C.P., J.S.W., and L.A.G. have filed patent applications on the platform/materials presented in this report. J.S.W. serves as an advisor to and/or has equity in 5 AM Ventures, Amgen, Chroma Medicine, KSQ Therapeutics, Maze Therapeutics, Tenaya Therapeutics and Tessera Therapeutics. L.A.G. has filed patents on CRISPR tools and CRISPR functional genomics, is a co-founder of Chroma Medicine, and a consultant for Chroma Medicine. The other authors declare no competing interests.

## Methods

### Library design

We designed two libraries that differ in size. The small-scale library (L1) is comprised of 37 domains; 32 domains were extracted in an unbiased fashion from 20 genes and 5 domains come from previously characterized gene modulators^6,7,13,19^. The 20 genes represented in the small-scale library were selected as genes whose products modify gene expression via epigenetic or transcriptional modulation.

The large-scale library comprises 347 domains. 266 of these domains were extracted from 194 genes obtained from dbEM, a database of epigenetic modifiers^31^. 81 previously published domains were included as positive controls (Supplementary Table 1). The same domains of the small-scale library were included in the large-scale library (L2). The domains in the large-scale library range between 135 and 3861 base pairs, with a median length of 1301 base pairs. All library elements and domain IDs are included in Supplementary Table 3.

### Library cloning and amplification

Each N- and C-terminal fragment was synthesized in an arrayed fashion and pooled for combinatorial assembly (Twist Bioscience). Each N- and C-domain combination is fused via flexible linkers on either terminus of dCas9, i.e., N_DOMAIN-XTEN16-NLS-NLS-dCas9-XTEN80-C_Domain.

Our vector is driven by a SFFV promoter with the following general structure: SFFV-N_DOMAIN-XTEN16-NLS-NLS-dCas9-XTEN80-C_Domain-IRES-PURO.

For ease of cloning, N-position domain fragments have homology with 3’ side of the SFFV promoter and the 5’ side of our dCas9 constant region (spanning XTEN16 through to the XTEN80), and the C-position domain has homology with its flanking XTEN80 and IRES sequences. Of the 347 domains in the library, ∼95% were synthesized successfully, resulting in a total of 110,554 dual terminus dCas9 fusion proteins.

The library was amplified using competent Stellar *E. coli* cells (Takara) and the library was isolated using the ZymoPure II Gigaprep kit (Zymo) following manufacturer’s instructions.

### Cell culture

K562 cells were cultured in RPMI-1640 supplemented with 10% FBS (VWR) and 1% penicillin, streptomycin, glutamine (Gibco) (termed RPMI-complete). K562 cells in static culture were maintained at 37°C and 5% CO_2_. When K562 cells were grown in shaking culture, 1X Pluronic F-68 (Gibco, Cat. 24040-032) was added to RPMI-complete media to reduce shear stress. K562 cells in shaking culture were maintained at 37°C, 8% CO_2_, and a shaking rate of 120 rpm in a humidified shaking incubator. Cryopreserved cell stocks were made with 10% DMSO in FBS (VWR). All cells in culture were tested and free of Mycoplasma (Lonza, cat. LT07-710).

### K562 cell line generation

K562 cells were transduced with a single guide RNA targeting *CD55* (g56) and sorted on BFP expression on a BD FACS Aria Fusion, resulting in K562-g56 cells. Validation of effector constructs were carried out in K562s expressing gRNAs per target gene using either CRISPRa or CRISPRi-designed gRNAs for activating or repressive constructs, respectively. For dual-guide experiments targeting *CD55*, K562s were transduced with CRISPRi or CRISPRa-designed gRNAs (pCW5 and pCW6, respectively). For gene activation experiments, K562s expressing dual gRNAs targeting *CXCR4* and *IL-2* were generated via lentiviral transduction of pCW57 and pCW13, respectively. Cells were transduced in media supplemented with 8µg/mL polybrene and spun at 1000 x g for 30 minutes to 2 hours. After transduction, cells were sorted on a BD FACS Aria Fusion for BFP expression. See Supplementary Table 2 for guide sequence and construct information.

### K562 suspension cell screens

#### Small-scale library screen (L1)

K562 cells stably expressing a CD55 targeting sgRNA (g56) were first transduced with lentiviral particles encoding the construct library. Transduction was performed by centrifuging cells in 100% viral supernatant supplemented with 8 ug/mL polybrene at 1000g for 2 hours at 33°C. Following spinfection, cells were resuspended in fresh media and cultured overnight. The following day (day 1), infected cells were split into replicates and expanded for two days. Cells were stained for infection efficiency 2 days post-infection by intracellular dCas9 stain. Four days post infection, cells were harvested, resuspended in freshly prepared FACS buffer (1% BSA in DPBS) to a final concentration of 10e6 cells/ml, and stained with anti-CD55-APC (Biolegend) antibody for 25 minutes on ice. Stained cells were pelleted then resuspended in FACS buffer to a concentration of 10e6 cells/mL. Live cells were analyzed for CD55 expression on a BD FACSAria Fusion and ∼20e6 cells were sorted into low and high quartile bins. Unsorted cells were also harvested. Cell fractions were centrifuged at 800xg then cell pellets were frozen at -80°C.

#### Large-scale library screen (L2)

K562 cells stably expressing a CD55 targeting sgRNA were first transduced with lentivirus as done for L1 in K562 cells. Following spinfection, cells were resuspended in fresh media and the following day, infected cells were split into replicates. Two days following infection, cells were selected with 3 ug/mL puromycin in complete media. The following day, 3 ug/mL fresh puromycin was added to the media. Four days post infection, cells were resuspended in fresh media for recovery from puromycin treatment for 24 hours. The following day, cells were cyropreserved using standard procedure due to timing constraints. Cells were thawed using standard procedure and allowed to recover for three days before performing the sort.

Eight days post infection and three days post thaw, cells were harvested, resuspended in freshly prepared FACS buffer to a final concentration of 10e6 cells/ml, and stained with anti-CD55-APC (Biolegend) antibody for 25 minutes on ice. Stained cells were pelleted then resuspended in FACS buffer to a concentration of 10e6 cells/mL. Live cells were analyzed for CD55 expression on a BD FACSAria Fusion and ∼40e6 cells were sorted into low and high quartile bins, corresponding to >400-fold coverage of the library. All cell fractions were centrifuged at 800xg and cell pellets were frozen at -80°C.

### iPSC Cell Culture and Screening

#### Cell line maintenance

Passage 4 KOLF2.1J iPSCs (male) were obtained from Jackson Laboratories and used for subsequent iPSC cell line generation. Briefly, cells were thawed onto Cultrex (R&D Systems, Cat. 3434-010-02) coated plates in mTeSR Plus complete media (Stem Cell Technologies) supplemented with 1 μM thiazovivin (THZ) (Sigma-Aldrich, cat. SML1045-25MG). Routine passaging was done in mTeSR plus media containing THZ. 1-2 days post-passage cells were maintained in mTeSR Plus media without THZ supplement. Cells stocks were frozen in 200ul of Synthafreeze (Thermo Fisher, cat. A12542-01).

#### Cell line generation

gRNA-expressing cells were generated using lentivirus-packaged gRNA plasmid (pCW9). KOLF2.1J wildtype cells were transduced with 200ul lentivirus in 800ul mTeSR Plus and THZ. 1 day post-transduction, media was changed to wash out lentivirus. 2 days post-transduction, cells were harvested using StemPro Accutase (Gibco, cat. A1110501) to check BFP expression levels (indicative of successful construct integration). Cells were sorted on BFP expression on a BD FACS Aria Fusion and replated in mTeSR Plus supplemented with Antibiotic-Antimycotic (Gibco, cat. 15240096) and CloneR (STEMCELL Technologies, cat. 05888). After recovering from the sort, cells (now termed KOLF2.1J-PLV-LVpCW9) were cultured in mTeSR Plus without CloneR or Antibiotic-Antimycotic. All cells in culture were tested and free of Mycoplasma (Lonza, cat. LT07-710).

#### iPSC Library 1 Screening

KOLF2.1J-PLV-LVpCW9 cells were plated to account for 1000X coverage of L1 and low multiplicity of infection library integration. All iPSC counting was performed on a Nexcelom Cellometer Auto 2000 using acridine orange/propidium iodide (AOPI) stain (Nexcelom). (T0) Approximately 12.8×10^6^ cells were plated on cultrex-coated T175s, in 20mL mTeSR Plus and 1µM THZ, so that the cells cover the surface area of the plate in low volume (as previously described^48^). 5mL of virus was added to each flask and the cells and virus sat at room temperature in the BSC for 10 minutes. The cells were moved to the incubator and 6 hours later, additional media was added on top of the attached cells (25 mLs of prewarmed mTeSR Plus + THZ). (T1) The following day, a complete media change was done with pre-warmed mTeSR Plus. (T2) Cells were harvested and split onto 4 T175s in mTeSR Plus and THZ containing 35mL media per flask. 1e6 cells per flask were taken for Cas9 intracellular staining to assess library integration. (T3-T6) Media changes were completed daily with fresh puromycin: two days of 0.8µg/mL and two days of 1µg/mL in mTeSR Plus. (T7) Media was replaced with pre-warmed mTeSR containing no puromycin. (T9) Cells were harvested for cell sorting.

Briefly, each flask was aspirated and 12mL accutase was added to each flask. Cells were incubated with accutase for 5-8 minutes and then cell suspensions were harvested for centrifugation at 800rpm for 4 minutes. Then, cells were resuspended in mTeSR Plus + THZ and counted using AOPI. Cells were harvested prior to cell staining for the unsorted condition per replicate and then the remaining cells were taken for CD55 staining and FACS. Cell pellets for each bin (top and bottom 20-25% CD55 expression) were harvested, pelleted, aspirated, and stored at -80°C prior to genomic DNA isolations.

### dCas9 Intracellular staining

Cells transduced with COMET library (and control cells) were pelleted by centrifugation and the supernatant was aspirated. Cells were resuspended in approximately 100 µl 4% formaldehyde (Thermo Scientific, cat. 28906) per 1 million cells and mixed well. Cells were fixed for 15 min at room temperature. Cells were washed with excess 1X PBS and centrifuged, followed by resuspension in 0.1 ml 1X PBS (cells may be stored overnight at 4°C in PBS at this point). Cells were permeabilized with 900µl ice-cold 100% methanol (resulting in 90% methanol final concentration) and left on ice for 10 minutes. Cells were washed with excess PBS, centrifuged, and the supernatant was discarded. The wash step was repeated 2 more times. Cells were resuspended in 100ul diluted conjugated Cas9 antibody (Cell Signaling Technologies, cat 7A9-3A3) in FACS Buffer (1X PBS with 1 % BSA) and incubated at room temperature in darkness for 1 hour. Cells were washed with FACS buffer 2X and spun at 200g for 5 minutes between every wash. Cells were analyzed on an Attune NxT Flow Cytometer.

### Library preparation

Genomic DNA was isolated using Macherey Nagel Blood or Blood XL kits following manufacturer’s instructions. Lentivirally integrated library constructs were amplified by PCR in 100ul reactions (5µg gDNA/reaction) using NEBNext Ultra II Q5 Master Mix (NEB Cat. M0544) from genomic DNA using dual-barcoded primers and the following PCR protocol: 98°C for 30s; 28 cycles x [98°C for 10s, 71°C for 30s, 72°C for 11min]; 72°C for 5min; hold at 4°C. PCR primers included barcodes for multiplexing and pooling prior to SMRTBell library preparation (Supplementary Table 8).

PCR cleanup was done using 0.45X AMPure PB beads (Pacific Biosciences). Briefly, AMPure PB beads were incubated with PCR product for 30 minutes at room temperature. PCR-bead mix was added to a magnet, washed 2X with 80% EtOH, and eluted in 40µl of Elution Buffer (Pacific Biosciences). Library size selection for amplicons between 4.5 and 14kb was conducted using a BluePippin (Sage Science) with 0.75% cassettes (Cat. BLF7510) and the 0.75% high pass 6kb-10kb v3 run protocol. Size-selected amplicons were removed from the elution well of the BluePippin cassette 45 minutes after elution finished, and wells were washed 2X with 40µl of 0.1% Tween20. Following size selection, amplicons were cleaned up using 1X SMRTBell cleanup beads (See manufacturer’s instructions, Alternate Size Selection Methods) and eluted in 10µl of Elution Buffer (Pacific Biosciences). Size-selected amplicons were prepared for sequencing using the SMRTBell Library Prep Kit 3.0 (Pacific Biosciences) following manufacturer’s instructions. Concentration was measured using Qubit dsDNA HS (Thermo Cat. Q32851) and size was measured on an Agilent Tapestation 4200 using Genomic DNA ScreenTapes or using a Bioanalyzer (Agilent). Replicates & conditions were pooled in an equimolar manner.

### Long read sequencing and data analysis

Amplicon libraries were sequenced on Sequel IIe and Revio systems (Pacific Biosciences).

The workflow and code used for long read sequencing processing is available at https://github.com/cmwilson24/Domain_Screening. PacBio library alignment code is available at https://github.com/GilbertLabUCSF/dCas9-fusions.

#### Circular consensus sequence (CCS) generation

subreads.bam files were exported from Sequel IIe (Pacific Biosciences) and Circular Consensus Sequences (CCS) were generated using ccs^49^ to produce a BAM file. Default parameters were used with the addition of the threads parameter (-j 40). Revio high-fidelity (HiFi) reads were automatically processed to BAM files.

#### Demultiplexing of BAM files

BAM files of CCS were demultiplexed using lima (Pacific Biosciences, https://github.com/PacificBiosciences/pbbioconda) to produce BAM files for each sample. Default parameters were used with the addition of the threads parameter (-j 10).

#### Sequence processing

Briefly, individual BAM files were converted to fastq files using bam2fq (samtools^50^) and trimmed using trimfq (https://github.com/lh3/seqtk) such that 16 nucleotides on either end were excluded. The parameters were as follows: seqtk trimfq -b 16 -e 16 sample.ccs.fq > sample.ccs.fq.

After trimming, samples were processed using scripts developed based on *knock-knock*^51^ to generate three-part alignment counts (Supplementary Figure 1A). Implementation of our alignment pipeline is available here: https://github.com/GilbertLabUCSF/dCas9-fusions. Briefly, a count threshold of 10 was set for all domain pairs per replicate unless otherwise described. For replicate correlations, the fraction is calculated by dividing the counts per library combination by the total reads per sample (condition and replicate). Following count filtering, DESeq2^26^ was used for downstream analyses with condition (high vs low expression) used as the covariate (“Design”). DESeq2 produced Log_2_Fold Change (Log_2_FC) scores comparing domains enriched in the high- and low-expressing bins and associated p-values.

### Epistasis analysis

To estimate the contribution of each domain individually to the overall change in gene expression, we fit a least-squares regression to estimate the Log_2_FC for each individual domain. To account for potential position-specific effects, we estimate separate Log_2_FC values for the N- and C-terminal versions of each domain. We first select domains with a count threshold of at least 50 for each biological replicate. If dA_N_B_C_ is the Log_2_FC of the paired perturbation

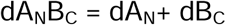

We observe that most combinations are well predicted by this model (Supplementary Figure 3A), confirming our assumption that most pairs are non-interacting. We then examine domain pairs whose predicted expression under this model differs substantially from the actual observed expression. When such a difference arises in both biological replicates, we view this as evidence supporting an interaction.

### Arrayed construct cloning

Arrayed validation hits were cloned by PCR amplification of individual N and C-terminal fragments, the constant XTEN16-NLS-NLS-dCas9-XTEN80 fragment, and vector backbone (pCW4). We used a modified destination vector for arrayed cloning with an added P2A-mNeonGreen protein after our IRES-PURO elements. All reactions were digested with Dpn1 (NEB) for at least 30 minutes prior to cleanup. PCR products were run on 1% agarose gel stained with 1X SYBR-Safe (Thermo) and extracted using the NucleoSpin Gel and PCR Cleanup Kit (Macherey-Nagel) according to manufacturer’s instructions. Constructs were assembled by Gibson Cloning using 2X HiFi DNA Assembly Master Mix (NEB) and transformed in Stellar *E. coli* cells (Takara). Colonies were isolated and miniprepped using either QIAprep Spin Miniprep Kit (Qiagen) or NucleoSpin 8 Plasmid Core Kit (Macherey-Nagel) and sequenced via whole plasmid sequencing prior to downstream assays.

### Lentivirus production

Library lentivirus was generated using LV-MAX Lentiviral Production System (Thermo, cat. A35684) per manufacturer’s instructions. LV-MAX cells were subcultured in LV-MAX serum-free production medium after reaching > 95% viability. Suspension cells were transfected in 2L flasks and 5-6 hours later, LV-MAX Transfection enhancer was added. 2 days later, liquid culture was spun at 1300 x g for 15 minutes. Supernatant was harvested and filtered through 0.45 µm filters. Aliquoted virus was frozen on dry ice and then stored at -80°C.

Lentivirus for validation constructs were generated using Lenti-X (Takara) or HEK293T cells in 6 well plates in 3mL DMEM supplemented with 10% FBS (VWR) and Pen-Strep (Gibco). Briefly, 400-600K cells were seeded onto well so 6 well plates the day before transfection. Per condition, 1.5µg transfer vector, 1.35 µg of packaging vector pCMV-dR8.91, 165 ng of packaging vector pMD2-G were combined into 7.5uL of TransIT-LT1 (Mirus Bio, MIR 2300) in 300uL Opti-MEM (Gibco). ∼16 hours later, media was replaced with fresh DMEM-complete and 1X ViralBoost reagent (ALSTEM Bio). Two days later, viral supernatant was harvested and filtered using 0.45µm filters.

### Arrayed K562 validation

Validation constructs were screened for activator or repressor activity by lentiviral integration, with the addition of negative control (scrambled) fusion protein (pCW31). Briefly, ∼200k cells expressing dual-gRNAs for target genes of interest (or single guide for epistasis analyses and screen phenotype comparisons at *CD55*) were transduced with 200ul of lentivirus in 2 replicates in a total volume of 400ul (RPMI complete + 8µg/mL polybrene transfection reagent (Millipore)) per well of a 48-well plate. Cells were spun for 2 hours at 1000g. After spinfection, cells were resuspended and additional media (∼200ul of RMPI-complete) was added per well. Cells were assayed by antibody staining and flow cytometry 3-9 days post-transduction.

### Antibody staining for flow cytometry

Cells were stained for flow cytometry as previously described^25^. Briefly, 200ul cells were harvested into 96-well plates. All centrifugation steps were performed at 500g for 5 minutes at 4°C followed by decanting of supernatant. Cells were spun down and washed 1x with FACS Buffer (PBS with 1% BSA). Then, cells were resuspended in 50ul FACS Buffer containing 1:100 dilution of antibody of interest and incubated at 4°C for 30 minutes. Cells were washed 2X with FACS Buffer and resuspended in 200ul of FACS Buffer. All flow cytometry was performed on Attune NxT instruments (Life Technologies). For knockdown/overexpression experiments, all data points shown in figures are events first gated in FlowJo for single cells based on FSC/SSC, then gated on BFP-positivity as a marker for stable gRNA expression, followed by gating on mNeonGreen-positivity as a marker for cells successfully transduced with fusion protein constructs.

Gated data was exported and analyzed in R using two functions from bears01 (https://github.com/chris-hsiung/bears01/tree/master/R): flowjo2df.R and calculate_quantilethresperc.R. All plots were generated in R.

### qPCR

K562s expressing pCW13 (IL-2 CRISPRa guides) were transduced with effector constructs (as described above). On day 5 post transduction, ∼100k cells harvested and washed with PBS prior to storing as pellets at -80°C. Cells were thawed on ice and lysed using the *Power* Sybr Green Cells-to-Ct Kit (Thermo). Reverse transcription and RT-PCR were performed per kit instructions and qPCR reactions were performed in triplicate on a Roche LightCycler 480. IL-2 primers were nominated using IDT’s PrimerQuest tool.

GAPDH Primers:

GAPDH-F: GGAAGGTGAAGGTCGGAGTC
GAPDH-R: GTTGAGGTCAATGAAGGGGTC

IL-2 Primers:

IL2-F: CTCCAGAGGTTTGAGTTCTTCT
IL2-R: AAACTCACCAGGATGCTCAC

### CUT&Tag

100K cells + 10% overage were harvested per condition, with two biological replicates. Briefly, 110k cells were harvested per antibody and effector condition and nuclei were isolated according to EpiCypher’s CUTANA Cut and Tag kit. Isolated nuclei (confirmed via manufacturer’s protocol quality control steps) were split up per antibody reaction and we proceeded per manufacturer’s instructions. Samples were barcoded during PCR using varying i5 and i7 primers. Libraries were QC’ed on an Agilent Tape Station using DS1000 High sensitivity tapes and pooled accordingly. Samples were run on a NextSeq (Illumina) with PE 2×50 sequencing and demultiplexed on instrument.

Libraries were analyzed using the Nextflow nf-core/cutandrun pipeline^52^, with the following modifications:

nextflow run nf-core/cutandrun -r 3.2.1 --input ./samplesheet.csv --outdir ./results_norm -- genome GRCh37 --normalisation_mode CPM --peakcaller MACS2 -profile conda

Bigwig files for H3K27ac, H3K27me3, and H3K9me1 per effector condition were visualized in Integrative Genomics Viewer (IGV v. 2.16.2).

Many plot color palettes were selected from Bang Wong’s coloring for color-blind individuals^53^.

Further information and requests for constructs, data availability or resources should be directed to the lead contact, Luke Gilbert.

