## Supplementary material for "Combinatorial effector targeting (COMET) for transcriptional modulation and locus-specific biochemistry": Document S1

**Supplementary Figures S1-5.**

**Supplementary Figure S1.**

(A) Long-read sequencing library alignment schema. Circular consensus sequencing (ccs) is performed if samples were sequenced on a Sequel IIe, otherwise a Revio generates HiFi reads automatically. Samples are demultiplexed using lima (Pacific Biosciences) through the use of barcodes appended during PCR and reads are aligned to the library reference. Counts per domain combination are generated per sample.

(B) Replicate correlations per condition (CD55 high, low, unsorted, respectively), showing log_10_ normalized counts (count per domain combination/total counts per sample = fraction).

(C) Replicate correlations comparing domain combination enrichment in high relative to low CD55 expression bins, showing log2(high:low) per replicate. Points are colored by fusion protein length (bp). Here, high:low comparisons use normalized counts after setting a count threshold of 10 per condition.

**Supplementary Figure S2.**

(A) Replicate correlations comparing domain combination enrichment in high relative to low CD55 expression bins, showing log2(high:low) per replicate for L2 in K562 cells. Points are colored in red if the effector combination contains VP64 and in blue if the effector combination contains KRAB. Here, high:low comparisons use normalized counts after setting a count threshold of 10 per condition.

(B) Large format of Figure 2C. Heatmap of Log_2_FC (High:Low CD55 expression bins) for individual N-C effector combinations with log2FC values from DESeq2. N domains are on the y axis and C domains are on the x axis, with positive log2FC in blue (activators) and negative log2FC values in red (repressors).

(C) Median Log2Fold Change of CXCR4 expression as measured by flow cytometry in K562 cells expressing a dual-guide RNA targeting CXCR4 and normalized to the median CXCR4 expression of cells targeted with an inactive dCas9 fusion protein (pCW31). Here, guide-expressing K562 cells are transduced with combinatorial activation constructs and CXCR4 expression is assayed 6 days post-transduction.

**Supplementary Figure S3.**

(A) Top 20 most frequently observed significant (p-value < 0.05) N or C domains for activators (top, log2FC > 1) and repressors (bottom, log2FC < -1) from the L2 K562 screen at CD55, showing enriched N-terminus effectors in orange (left) and C-terminus effectors in pink (right).

(B) Upset plot categorizing N and C domain function for significant repressors (log2FC < -1 and p-value < 0.05) identified in the L2 K562 screen, showing a minimum intersection size ≥5.

(C) Upset plot categorizing N and C domain function for significant activators (log2FC > 1 and p-value < 0.05) from the L2 K562 screen with an intersection size ≥5.

**Supplementary Figure S4.**

(A) Scatter plot showing epistasis modeling predictions for effector combination activity on the x-axis for each replicate (Replicate 1, Left; Replicate 2, Right) and DESeq2 Log2FC values for effector combinations from the COMET L2 screen in K562 cells on the y-axis. The residual value per replicate is the distance between the expected and observed activity for a particular domain combination.

(B) Scatter plot showing residual values calculated from epistasis modeling using the L2 screen in K562 cells, showing Replicate 1 on the x-axis and Replicate 2 on the y-axis.

**Supplementary Figure S5.**

(A) CUT&Tag experiments detecting H3K27ac were repeated to reconfirm initial observations (Figure 5B & C). Traces visualized in IGV detecting H3K27ac marks in cells expressing a gRNA targeting *CD55* for two replicates per condition. Traces show H3K27ac in cells transduced with MBD2-CDYL (green) relative to parental K562 cells (black), with a decrease in H3K27ac in MBD2-CDYL treated cells.

(B) H3K9me1 CUT&Tag data for parental K562 cells (black), or cells transduced with MBD2-single (pink), single-CDYL (light blue), and MBD2-CDYL (green) effectors across two technical replicates as visualized in IGV across a 67kb region surrounding the *CD55* locus. We observe no appreciable differences in H3K9me1 across conditions.

(C) H3K27me3 CUT&Tag data for parental K562 cells (black) or cells transduced with MBD2-CDYL (green) effectors across two technical replicates. We observe no H3K27me3 repressive marks across the locus in either parental or treated conditions, with K562 ENCODE ChIPseq data included for reference.
