## Supplementary figures and images for "Combinatorial effector targeting (COMET) for transcriptional modulation and locus-specific biochemistry"

### Supplementary Figure 1

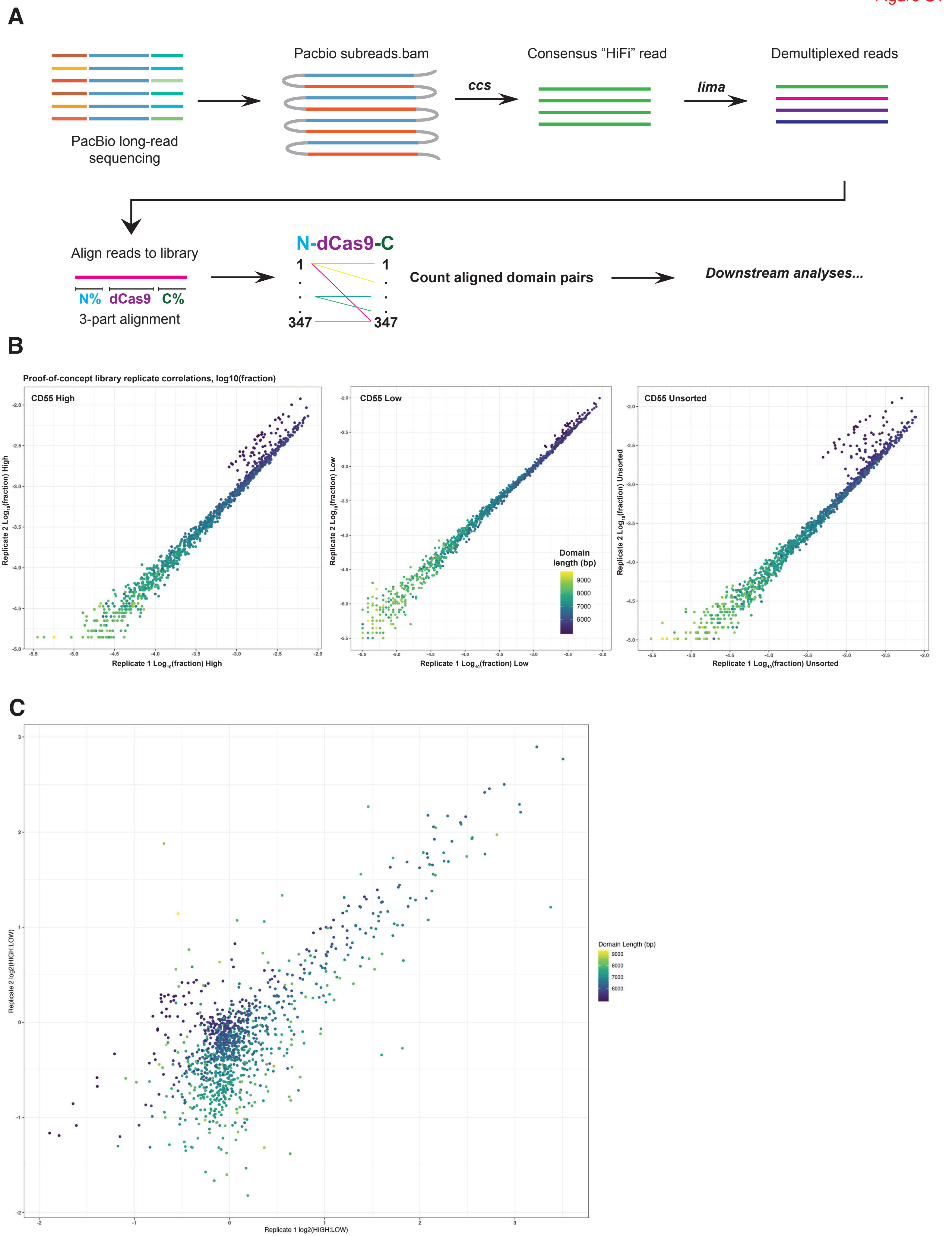

### Supplementary Figure 2

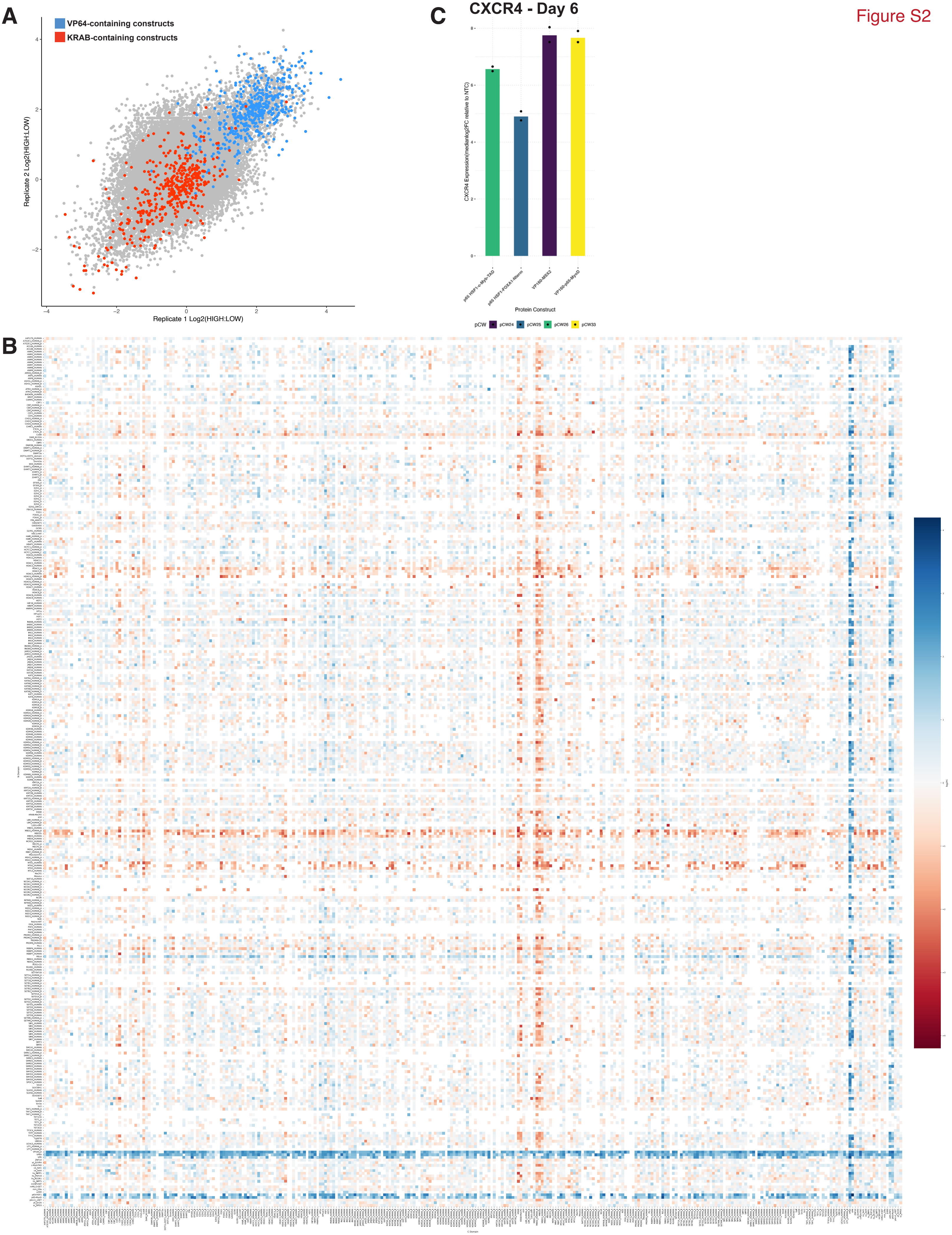

### Supplementary Figure 3

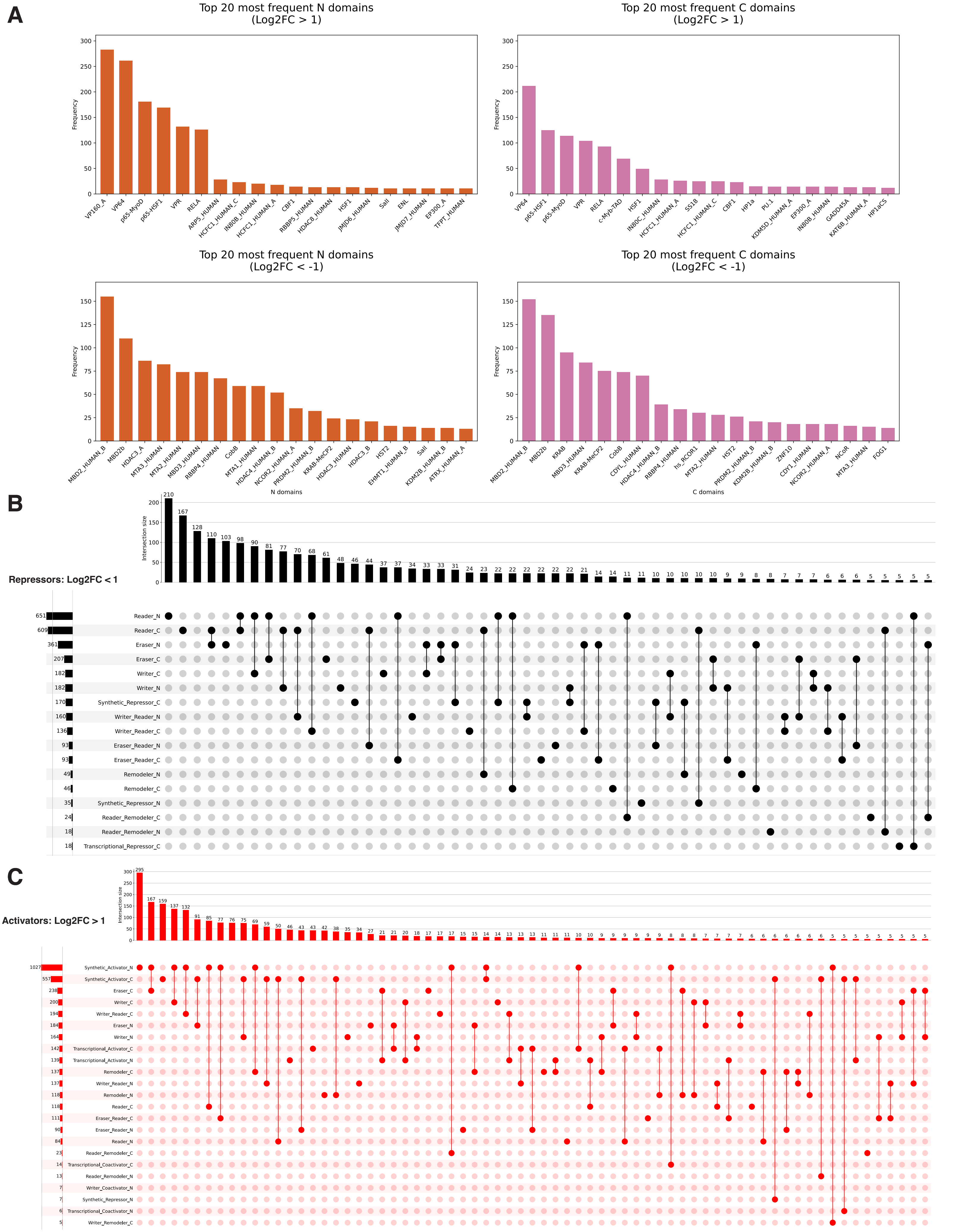

### Supplementary Figure 4

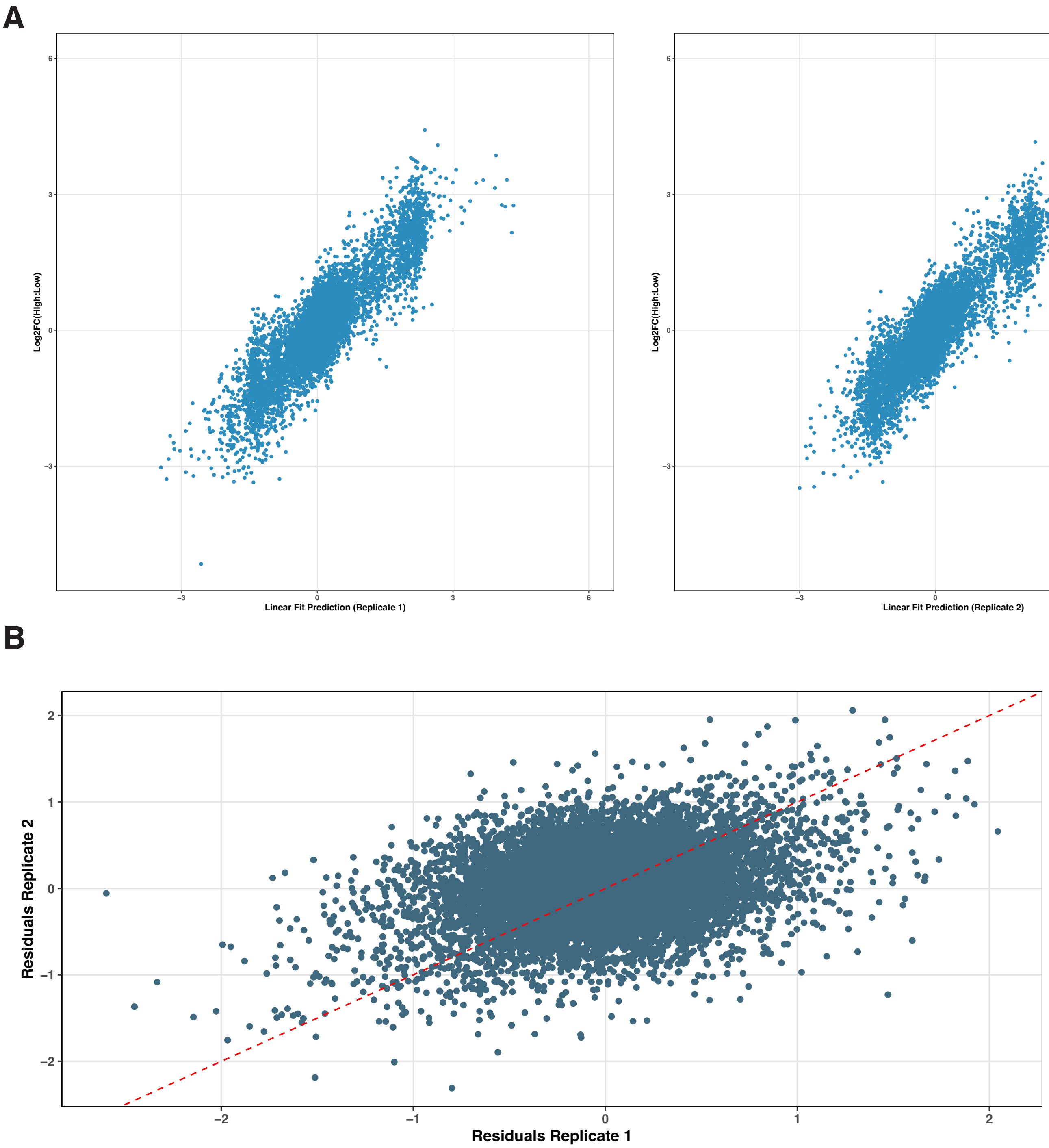

### Supplementary Figure 5

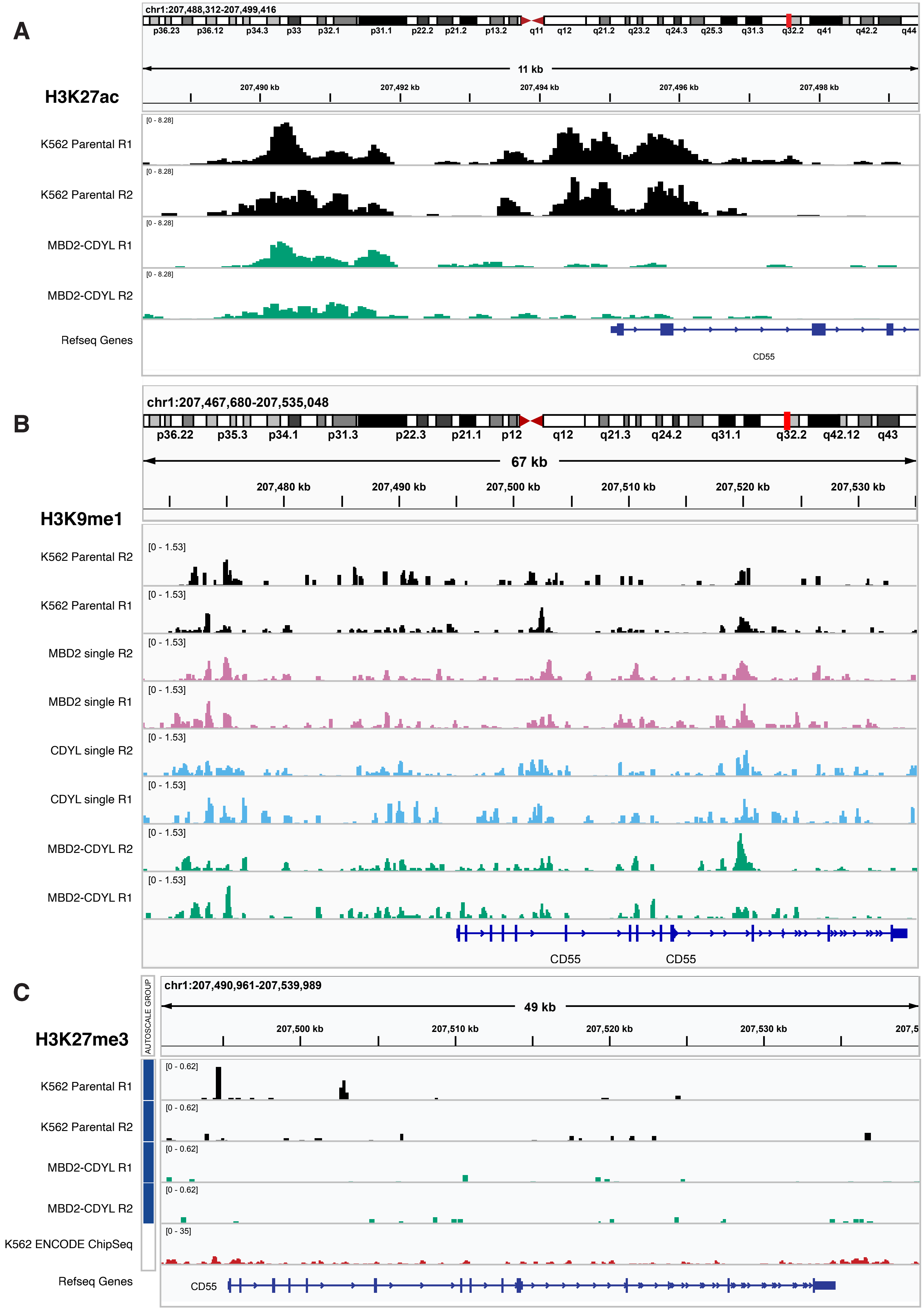
